# Efficient training of neural networks using natural vectors with covariates for the plant microRNA precursor prediction

**DOI:** 10.64898/2026.01.08.693543

**Authors:** Javier Montalvo-Arredondo, Marco Adán Juárez-Verdayes

## Abstract

The Fabaceae plants (Legumes) are important for the economy and food sovereignty of México. Traits development of agronomic interest, and other biological activities of the Fabaceae plants are tightly related to the gene regulation, like the post-transcriptional gene repression mediated by microRNAs. Several artificial intelligence models have been developed for the miRNA precursor sequence prediction. They were based mainly on Convolutional Neural Network and Multi-Layer Perceptron architectures. Although the numerical encoding of nucleotide sequence and its secondary structure of pre-miRNAs implemented in these neural networks showed good performance, there are other encoding methods that have not been explored. Recently, a geometric construction of viral genome space and the numerical encoding of the archaea, bacteria, fungi and viruses genomes were successfully achieved employing natural vectors with covariance component. Natural vectors have also been used as input data during neural networks training for the classification of viral genomes. In consequence, in this work we mainly assessed the performance of neural networks as regression or classifier models trained with nucleotide sequences and its secondary structure representation encoded by natural vectors with covariance component alone or nested within the three sequences method. Additionally, we tested other characteristics of neural networks, and the results of training neural networks with natural vectors with covariates showed a better performance in predicting intrinsic nucleotide features, such as percentage of guanine and cytosine, pairwise-aligned sequence identity. Also, it showed good accuracy in categorizing miRNA precursor sequences compared with the results obtained from other encoding methods, that are often used in the numerical representation of nucleotide sequences.

## 1. INTRODUCTION

Some plants, which belongs to Fabaceae family, are of outmost importance for the economy and food sovereignty of México, there are more than five hundred of legume species reported in this country that are very useful, some of them are used for medicinal purposes, meanwhile, other species are cultivated to use them in animal and human food supply, it is because they are main sources of carbohydrates and essential amino acids, and the legumes farming activities in México are very intense and dynamic. (Baez-Mora et al., 2023; Centeno-González et al., 2021; Delgado-Salinas et al., 2021; Estrada-Castillón et al., 2024; Shavanov, 2021).

Traits development of agronomic interest, and other biological activities of the Fabaceae plants are tightly related to the gene regulation, like the post-transcriptional gene repression mediated by small RNA molecules of size ranging from 21 to 24 nucleotides called microRNAs (miRNAs) (Zhang et al., 2022b). Many scientific reports associate the miRNA regulatory system with the biotic and abiotic stress tolerance (Brant and Budak, 2018; Budak et al., 2015; Li et al., 2017; Shriram et al., 2016; Zhang et al., 2022a), the resistance to pathogen infection (Ouyang et al., 2014; Su et al., 2018; Yang et al., 2021; Yang et al., 2015; Zhu et al., 2013), the plant-environment interaction (Song et al., 2019), and the control of the development process (Ayubov et al., 2019; Dong et al., 2022; Feng et al., 2016; Li et al., 2017; Wu et al., 2009; Zhang et al, 2022b).

The regulatory complex associated with miRNAs is called RISC and is composed of a protein ensemble and the miRNA molecule that guides the complex to the mRNA targets. MiRNAs are encoded in protein and non-protein coding genes. The transcripts of these genes, which are called primary transcripts (pri-miRNAs), form a stable hairpin that is recognized and processed by the DICER-LIKE 1 endoribonuclease with the help of the Serrate and Hypnoatic leaves 1 cofactors to form the miRNA precursor (pre-miRNA). The pre-miRNAs are subsequently translocated to the cytoplasm, where they mature and form the RISC complex. (Gangadhar et al., 2021; Zhang et al., 2022b). To understand the gene regulation mediated by miRNAs in plants, it is important to correctly identify and locate the miRNA loci in the plants genome. That was a very difficult task to accomplish because the early main and ancillary criteria used for the miRNA genome annotation didn’t provide enough evidence (Kozomara and Griffiths-Jones, 2011; Meyers et al., 2008).

Currently, several artificial intelligence (AI) models have been developed, and they accurately predict miRNA precursor sequences and efficiently annotate miRNA encoding loci in plants. These AI models are based mainly on Convolutional Neural Network (CNN), whose sequence encoding methods are focused on the one-hot encoding with additional information derived from the secondary structure representation (Zhang et al., 2024; Zheng et al., 2019) and Multi-Layer Perceptron models (MLP) that use vectors composed of more than 180 features extracted from nucleotide sequences, secondary structure of the hairpin, and the thermodynamic properties (Lokuge et al., 2022), and vectors of n-grams (a.k.a vectors of k-mer frequencies or BoW vectors) that represent mainly the nucleotide sequence and the secondary structure of miRNA precursors (Juárez-Verdayes and Montalvo-Arredondo, 2025). Numerical representation (encoding) of nucleotide sequence and its secondary structure of pre-miRNAs is a crucial step in neural network training and prediction. Although several approaches have been proposed during the development of AI for the identification of miRNAs, other approaches remain unexplored for this purpose (Bohnsack et al., 2022).

Recently, a geometric construction of viral genome space was successfully achieved by means of finding the convex hull using the nucleotide sequence information encoded in natural vectors with covariance component. Also, this method was used for the numerical representation of the archaea, bacteria, fungi and viruses genomes for the taxonomic classification and phylogenetic studies (Sun et al., 2021; Sun et al., 2022). Additionally, they have been recently used in the training of neural network models for classifying viral genomes (Shi et al., 2025). In consequence, this method for the nucleotide sequence encoding has caught our attention, because we think it could be efficiently used for the encoding of pre-miRNA nucleotide sequence and its secondary structure for the neural network training. Accordingly, in this work, we assessed two encoding methods to pre-process the input data, namely, the natural vector with covariates (NV) and the natural vector with covariates nested within the three sequence method (TSM-NV). As a reference encoding method, we employed the bag of words vector (BoW, a.k.a n-grams or k-mer frequencies vector), and we also used the Frequency of the Chaos Game Representation (FCGR) as a method that only represents but doesn’t encode the nucleotide sequence (Bohnsack et al., 2022).

First, we evaluated the effect of these encoding methods for the nucleotide sequence pre-processing on the predictive performance of MLP regression and classifier models built for the prediction of guanine and cytosine percentage and the percentage of identity of pairwise sequence alignments. Additionally, we tested MLP or CNN models trained for miRNA precursor sequence classification trained with input data pre-processed by the encoding methods described above. We also assessed three activation functions, such as hyperbolic tangent (tanh), Rectified Linear Unit (relu) and leaky Rectified Linear Unit (leaky_relu); two manners of streaming the input data that are, one stream (1s; 1 channel), where different input data are concatenated, then passed to the neural network, or two streams (2s; 2 channels. a.k.a branched input), where different input data is handled independently by separated neural networks, then, the output is concatenated and canalized to a final neural network, similarly to the work of (Cha et al., 2021; Georgakilas et al., 2020); and two neural network architectures, such as MLP, mainly for the natural vectors, and CNN mainly for the case of the three sequence method nested within natural vectors that returns a matrix.

We obtained interesting results suggesting that encoding the neural network input data by the NV or TSM-NV method, is a streamlined option to encode the nucleotide sequence and its secondary structure representation for developing neural networks trained for the miRNA precursors prediction and intrinsic features of nucleotide sequences, obtaining competitive values of accuracy, sensitivity and specificity. Additionally, we observed a slight tendency to obtain better performance in models that were built with Rectified Linear Unit and leaky Rectified Linear Unit activation functions, and in some cases, we observed better performance in models where two input streams (2s; 2 channels) were implemented.

## 2 METHODS

### 2.1 Data acquisition

The nucleotide sequences used for the training and testing of the Multi-Layer Perceptron (MLP) models built for predicting the percentage of guanine and cytosine were randomly generated by computer using a Python3 custom script. The size of the sequences was varied between 60 to 200 nucleotides. We generated a dataset with 118,440 sequences for training, and a dataset with 29,610 sequences for testing.

To build the nucleotide sequence datasets for the training of MLP models for predicting the percentage of identity of the pairwise sequence alignment, we randomly select with replacement a pair of sequences from the previously described datasets, and the percentage of identity was calculated form local alignments performed with the Smith-Watermann algorithm with the parameters set as default (Smith and Waterman, 1981) implemented in the PairwiseAligner class from Biopython package. We generated two datasets containing entries composed of two sequences and the associated percentage of identity: 100,000 entries for training and 10,000 entries for testing purposes.

To train MLP models built for sequence classification based on the percentage of guanine and cytosine, we randomly generated new datasets of sequences, as described previously, for training and testing. We generated three datasets of sequences, each one containing sequences of sizes ranging from 60 to 200 nucleotides. One dataset consisted of sequences with a low level of %GC (0-33), another dataset was composed of sequences with a medium level of %GC (33-67), and the third dataset had sequences with a high level of %GC (67-100). We generated these datasets for the three categories with 12,427 sequences for each category, for the training process, and three datasets with 4,142 sequences for each category for model testing. To train and test MLP or Convolutional Neural Networks 2D (CNN2D) models built for the prediction of the Fabaceae pre-miRNAs, we used the same datasets described in (Juárez-Verdayes and Montalvo-Arredondo, 2025) where pre-miRNA sequences were downloaded from PmiREN2.0 database (Guo et al., 2022). The Python functions and classes we used to generate all these datasets of sequences are freely available at GitHub: https://github.com/exseivier/UAAAN-IA/.

### 2.2 Models and hyper-parameters

We built 24 models based on the MLP architecture, consisting of the input layer, three hidden dense layers of 500 neurons, and an output neuron. The input and output layers were varied to fit the input pre-processed data dimension, and the type of output was also varied (e. g. single neuron for regression or three neurons for classification tasks). We tested different types of input data encoding methods, namely, natural vector with covariates (NV), bag of words (BoW), vector of frequencies for the chaos game representation (FCGR), and the natural vector with covariates nested within the three sequences method (TSM-NV), which detailed description can be found in the sections below. We also tested different activation functions between hidden layers, such as hyperbolic tangent (tanh), no activation or linear activation (No act), Rectified Linear Unit (relu), and Leaky Rectified Linear Unit (leaky_relu). For those MLP neural networks built as regression models to predict the percentage of guanine and cytosine of a sequence, and to predict the percentage of identity, given a pairwise sequence alignment, we set one output neuron with an linear activation function; and for those neural networks built for categorizing sequences based on the %GC, we set three neurons as the output layer with the “softmax” activation function. For each model, we tested the four data input pre-processing (encoding) methods described above. We described in more detail the characteristics of the 24 models in (Table 1).

**Table 1.** Description of the MLP architectures trained for predicting the percentage of guanine and cytosine, and identity.

| Model ID<sup>a</sup> | Training | Activation functions | Output | Dropout |
| --- | --- | --- | --- | --- |
| 001 | Predicting %GC | Hyperbolic tangent. tanh(x) | 1 neuron, linear activation | NO |
| 002 | Predicting %GC | No activation function. | 1 neuron, linear activation | NO |
| 003 | Predicting %GC | Rectified linear unit. ReLU | 1 neuron, linear activation | NO |
| 004 | Predicting %GC | Leaky rectified linear unit. l-ReLU | 1 neuron, linear activation | NO |
| 005 | Predicting %GC | Hyperbolic tangent. tanh(x) | 1 neuron, linear activation | YES |
| 006 | Predicting %GC | No activation function. | 1 neuron, linear activation | YES |
| 007 | Predicting %GC | Rectified linear unit. ReLU | 1 neuron, linear activation | YES |
| 008 | Predicting %GC | Leaky rectified linear unit. l-ReLU | 1 neuron, linear activation | YES |
| 009 | Predicting % of identity | Hyperbolic tangent. tanh(x) | 1 neuron, linear activation | NO |
| 010 | Predicting % of identity | No activation function. | 1 neuron, linear activation | NO |
| 011 | Predicting % of identity | Rectified linear unit. ReLU | 1 neuron, linear activation | NO |
| 012 | Predicting % of identity | Leaky rectified linear unit. l-ReLU | 1 neuron, linear activation | NO |
| 013 | Predicting % of identity | Hyperbolic tangent. tanh(x) | 1 neuron, linear activation | YES |
| 014 | Predicting % of identity | No activation function. | 1 neuron, linear activation | YES |
| 015 | Predicting % of identity | Rectified linear unit. ReLU | 1 neuron, linear activation | YES |
| 016 | Predicting % of identity | Leaky rectified linear unit. l-ReLU | 1 neuron, linear activation | YES |
| 017 | Categorization task | Hyperbolic tangent. tanh(x) | 3 neurons, softmax | NO |
| 018 | Categorization task | No activation function. | 3 neurons, softmax | NO |
| 019 | Categorization task | Rectified linear unit. ReLU | 3 neurons, softmax | NO |
| 020 | Categorization task | Leaky rectified linear unit. l-ReLU | 3 neurons, softmax | NO |
| 021 | Categorization task | Hyperbolic tangent. tanh(x) | 3 neurons, softmax | YES |
| 022 | Categorization task | No activation function. | 3 neurons, softmax | YES |
| 023 | Categorization task | Rectified linear unit. ReLU | 3 neurons, softmax | YES |
| 024 | Categorization task | Leaky rectified linear unit. l-ReLU | 3 neurons, softmax | YES |

a. This model ID matches with the Python class name we used to build these neural networks.

### Table 1. Description of the MLP architectures trained for predicting the percentage of guanine and cytosine, and identity

We also built another 36 neural networks based on MLP or CNN2D architectures for the pre-miRNA sequences classification. With these models we evaluated different features, such as, (1) input data encoding methods (BoW, NV and TSM-NV), (2) input streaming, that means the input was either one stream (1s) where vectors, for MLPs, or matrices, for CNN2D, were concatenated first and then passed to the neural network, or two stream (2s) where vectors or matrices were passed to the model by independent neural networks composed by 3 hidden layers, and then the outputs were concatenated and canalized to an MLP neural network, and (3) two different architectures, MLP for BoW, NV and flattened TSM-NV vectors, and CNN2D for BoW and TSM-NV matrices. To construct the BoW matrices for CNN2D neural networks, we reshaped the BoW vectors computed from nucleotide sequences that yielded vectors of 1024 dimensions, whose reshaped matrix had a 32×32 dimension, and the BoW vectors computed from the pre-miRNA secondary structure sequence yielded vectors of 243 dimensions and were reshaped to matrices of 9×27 dimensions. For more information (see Table 2).

**Table 2.**
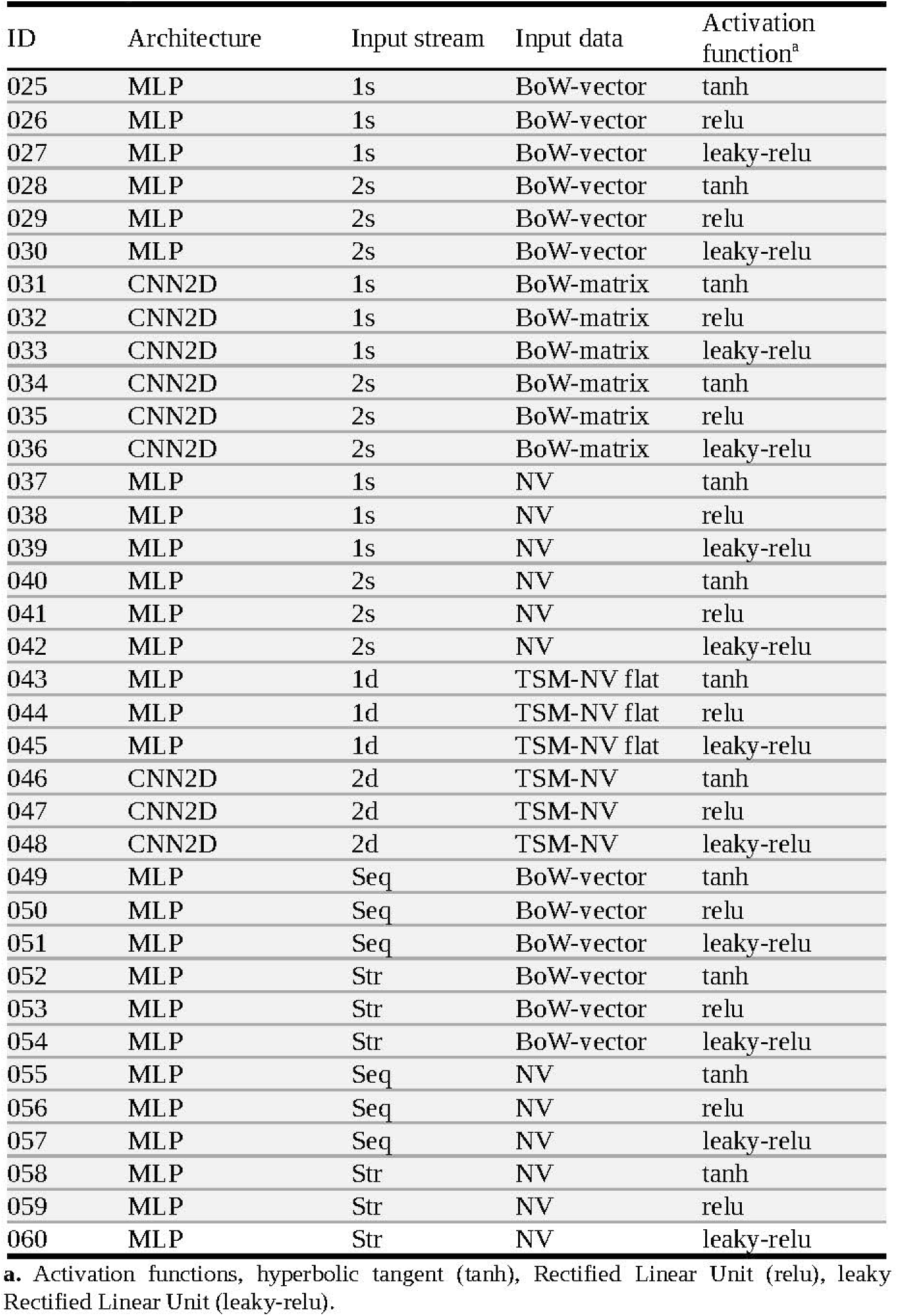
Description of the neural networks trained for predicting pre-miRNA sequences.

During the training of the neural networks used as regression models for the prediction of %GC and percentage of identity, we used the adaptive moment (Adam) as optimizer with a learning rate of 1×10 ^-3^ and a weight decay of 1×10^-6^. We used the mean squared error (MSE) as a loss function, and the training was done up to 1500 epochs using a data batch size of 64. For each epoch, the order of the dataset was randomized, and 80% of it was used for training, and 20% of it was used for validation. We monitored the progress of training and stored the loss values for the training dataset as well as for the validation dataset. We selected and saved the model that showed the lower validation loss value compared with the previously saved model. To train the MLP models used for categorizing sequences based on the %GC, we used almost all the parameters and methods described above, except for the loss function, which in this case was the categorical cross-entropy function, a common option for evaluating categorical output of artificial neural network predictions. For the case of the models trained for the pre-miRNA sequence classification task we used the same parameters except for the loss function and the activation function of the output neuron that were the binary cross-entropy and sigmoid functions respectively.

### 2.3 Input data pre-processing methods Natural vectors with co-variates

Natural vector (NV) represents the complexity of a DNA sequence in an 18-dimensional space composed of the frequencies of nucleotides, the average location distance, the *j*-th central moment of the nucleotide position, and the covariance between nucleotides (Bohnsack et al., 2022; Sun et al., 2021; Sun et al., 2022). And the formal definition is described as follows.

Given a DNA sequence *S*=*S*_1_*, S*_2_ *, S*_3_, … *, S_n_*, and an alphabet of the nucleotide letters *L*=**{ *A, C, G, T* }**. The indicator function is defined as,

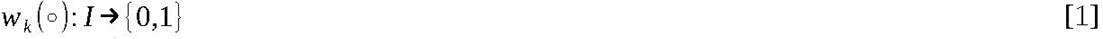

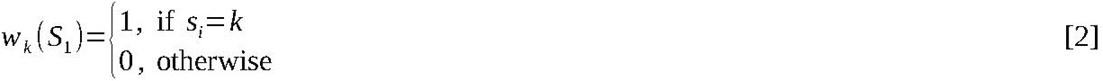

Where *s_i_, k* ∈ *L*, and *i*=1, 2,3, … *,n*. Then,

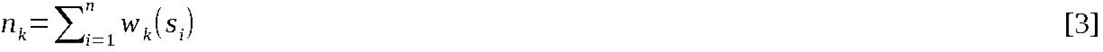

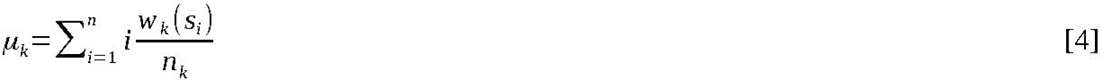

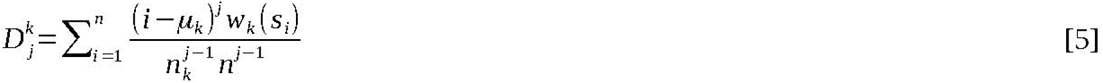

Where *n_k_* stands for the *k* nucleotide count, *μ_k_* is the average of the distance distribution for nucleotide *k*, and *D_j_^k^* is the *j*-th central moment for nucleotide *k*. To calculate the covariance for the nucleotide pair combinations, the indicator function is redefined as follows.

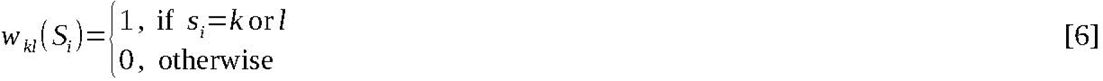

And the covariance is calculated with the following formula.

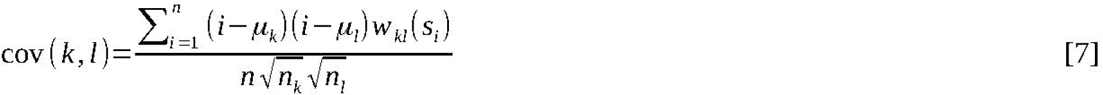

This pre-processing method returns an 18-dimansional vector with the following elements.

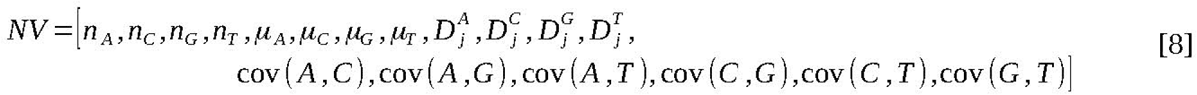

### K-mers counting and bag of words

The concept of bag of words (BoW) vector (a.k.a vector of k-mer/word frequencies or n-grams) is a set of unique feature frequencies (e. g. words) whose size depends on the total number of elements contained in the set. In biological sequences of DNA, RNA or amino acid residues, it is common to calculate the frequencies of the total possible kmers, where a kmer is a subsequence of size k for a given sequence (Martinc and Pollak, 2019).

The BoW vector can be defined as a dictionary or hash table *D* (*x*)→ *y* that maps x to y, where x is the kmer, and y is the frequency value of observing that kmer in a given sequence. This function can be used with the sliding window algorithm to extract the kmers of the sequence (a.k.a kmer decomposition) and calculate the frequencies for every possible kmer of size *k*.

In sliding window algorithm there are window boundaries defined by *i* and *i* +*k*, where *i* is the start and *k* is the kmer size. Given a sequence *S*={*s*_1_ *, s*_2_ *, s*_3_,… *, s_n_* }, where *s_i_* ∈ *L*={*A,C,G, T* } and a set of all possible kmers *M* ={*m*_1_ *, m*_2_ *,m*_3_,… *, m_p_* }, where *m_j_* can be represented as a subsequence of *S, m* =*S* [*s*_i_, …, *s*_*i*+*k*_] and *p*=len(*L*)*^k^*, that is the total number of possible kmers of size *k*, then a dictionary can be created as follows with a for loop.

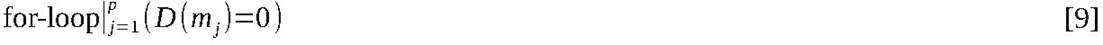

And the frequencies can be calculated using the kmer decomposition and the following for loop.

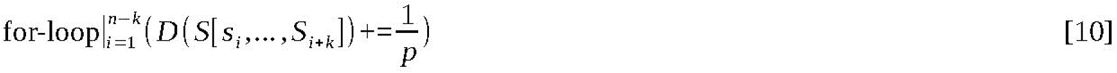

### Vector of frequencies for the chaos game representation

Chaos game representation is a pre-processing method that allows representing biological sequences, such as DNA or RNA, in a matrix of ℝ^2x^ *^n^* dimension. The elements of the sequence *S*={*s_1_, s_2_,* … *, s_n_* } are coded into vectors *x*_i_ ∈ℝ^2^ with the following formula (Löchel and Heider, 2021).

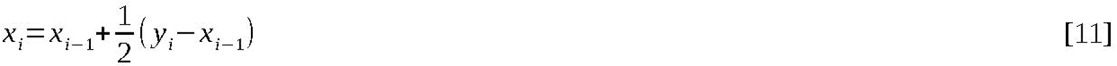

where *y_i_^T^*is defined as follows.

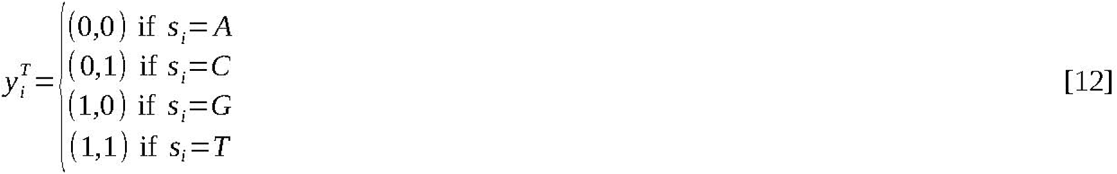

The calculation of the frequencies for the chaos game representation (FCGR) was achieved by the discretization of data calculated with [eq. 11], into a square matrix of defined size with the following for-loop.

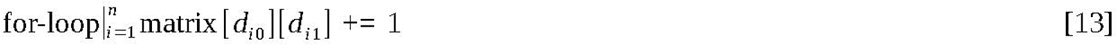

Where *d_i_* are the rounded values of the product of *x_i_*⋅*s*, and *s* is the size of one dimension for a given square matrix. To use this matrix in the feed-forward neural networks training and testing, the matrix was flattened.

### Three sequences method nested to natural vectors with covariance component

The sequences were pre-processed with the three sequences method (TSM) that returns a matrix of ℝ^3 × *n*^ dimensions where *n* is the length for a given sequence, where nucleotides were coded according to their properties based on the following indicator functions (Bohnsack et al., 2022).

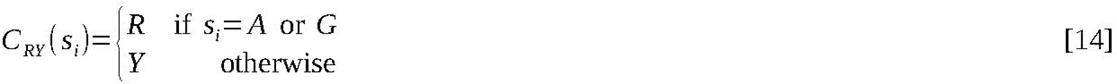

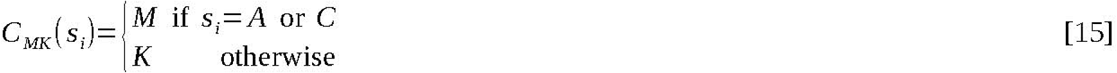

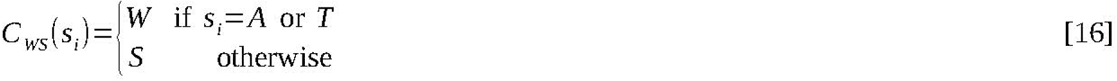

These indicator functions will render the matrix TSM∈ℝ^3 × *n*^.

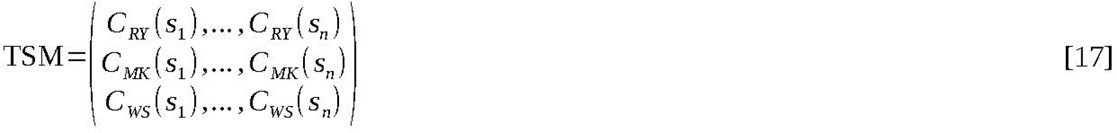

Every vector of the TSM matrix was processed with the NV method obtaining a matrix TSM-NV ∈ ℝ^3 × 18^, however, for training and testing MLP models, the matrix was flattened as described previously.

### 2.4 F1Score calculation

To measure the accuracy of mathematical models used in categorization tasks, it is usual to calculate the F1Score, which value represents a balanced estimation of the sensitivity and positive predictive value (PPV a.k.a Q-scores) scores for each category. The sensitivity is defined as the proportion of cases correctly predicted as true positives, for a defined category, from a population of known true positives, and the PPV is defined as the proportion of cases correctly identified as true positives for a category from a population predicted as true positives (Trevethan, 2017).

Based on these definitions, we calculated the sensitivity and PPV scores for the predictions made with MLP models trained for categorizing sequences based on %GC. These scores were calculated for every category (LOW, MEDIUM, HIGH; see materials and methods section). Given the confusion matrix shown in (**Table 3**).

**Table 3.** Confusion matrix for three categories.

|  |  | Predicted |  |  |
| --- | --- | --- | --- | --- |
|  |  | LOW | MED | HIGH |
| True | LOW | **TL** | FM<sup>1</sup> | FH<sup>1</sup> |
|  | MED | FL<sup>1</sup> | **TM** | FH<sup>2</sup> |
|  | HIGH | FL<sup>2</sup> | FM<sup>2</sup> | **TH** |

We calculated the sensitivity scores for every category as follow,

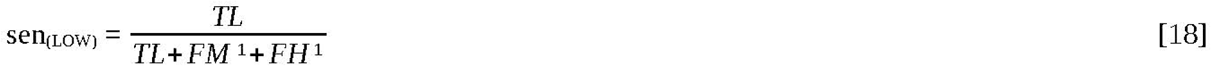

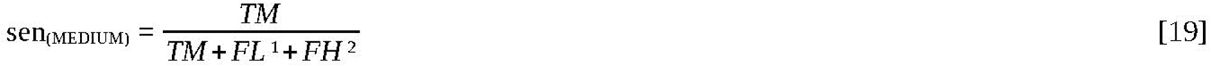

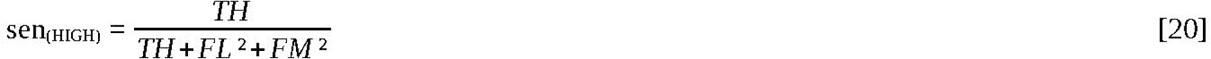

We calculated the PPV scores as follow,

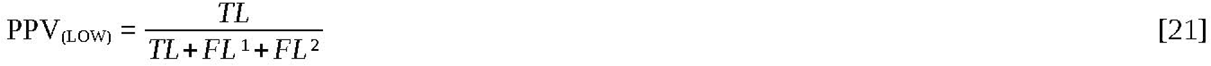

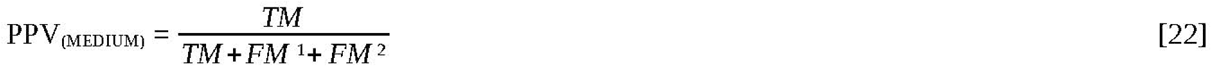

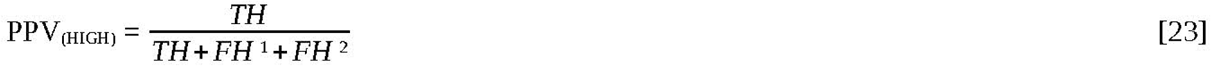

where TL, TM and TH stand for the LOW, MEDIUM and HIGH categories that were predicted as true categories. FL, FM and FH are these categories that were identified as false. And to calculate the F1Scores for the positive and negative predictions for every category, we used the following equations.

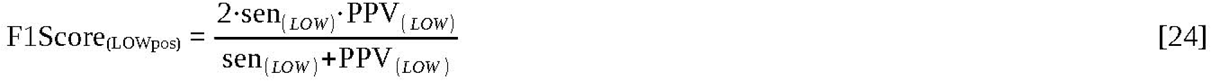

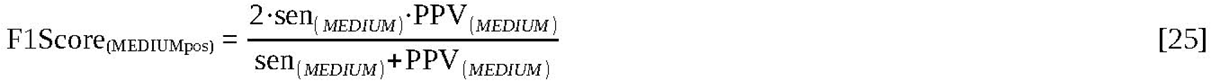

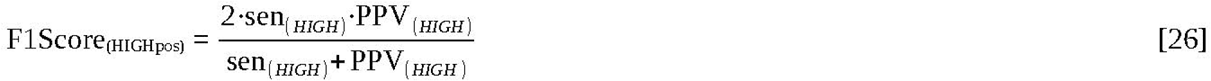

The log odds ratio values where calculated as follow,

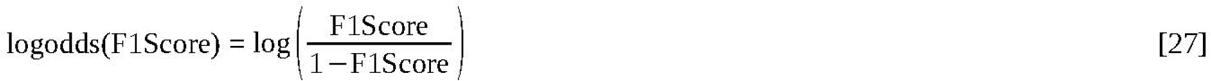

The code used to perform the encoding of nucleotide sequences and secondary structure representation, as well as to calculate the accuracy and precision values, is freely available at GitHub: https://github.com/exseivier/UAAAN-IA/.

## 3. RESULTS

### Predicting percentage of guanine and cytosine

In this analysis we built eight MLP models (001-008) configured with an input layer, three hidden dense layers of 500 neurons and an output neuron with a linear activation (No act). They were trained for predicting the percentage of guanine and cytosine for a given sequence data set, in which sequence size varied between 60 to 200 nucleotides. The first four models (001-004) were built with different activation functions set between the three hidden layers, namely, hyperbolic tangent (tanh), no activation function or linear activation (No act), rectified linear unit (ReLU), and leaky rectified linear unit (Leaky-ReLU). The last four models (005-008) were constructed in a similar way to the first four models, but in this case, dropout layers were set between hidden layers. We construct these models for each input data pre-processing method (BoW, FCGR, NV, TSM-NV).

To test the models’ predictive performance, we challenged them with the testing dataset and measured the mean squared error as loss value (MSE). We selected the loss value obtained from models trained with BoW vectors (MSE(*BoW*)) as the reference value, and calculated the log odds of the MSE(*X_i_*)/ MSE(*BoW*) quotient, where MSE(*X_i_*) is the loss value obtained from the models trained with the other encoding methods (*X_i_* ∈{FCGR, NV, TSM-NV}). In (**Figure 1**), we plotted the log odds MSE(*X_i_*)/ MSE(*BoW*) values in horizontal bars, and observed that models trained with NV method had a better performance (lower negative log odds values) compared with the worse results obtained with the FCGR method (higher positive log odds values) (**Figure 1A**). Despite of we observed that the predictive performance was negatively affected when dropout layers were used, the accuracy of the models trained with natural vectors remains higher than the accuracy observed for models trained with the FCGR and BoW input data encoding methods (**Figure 1B**).

**Figure 1.**
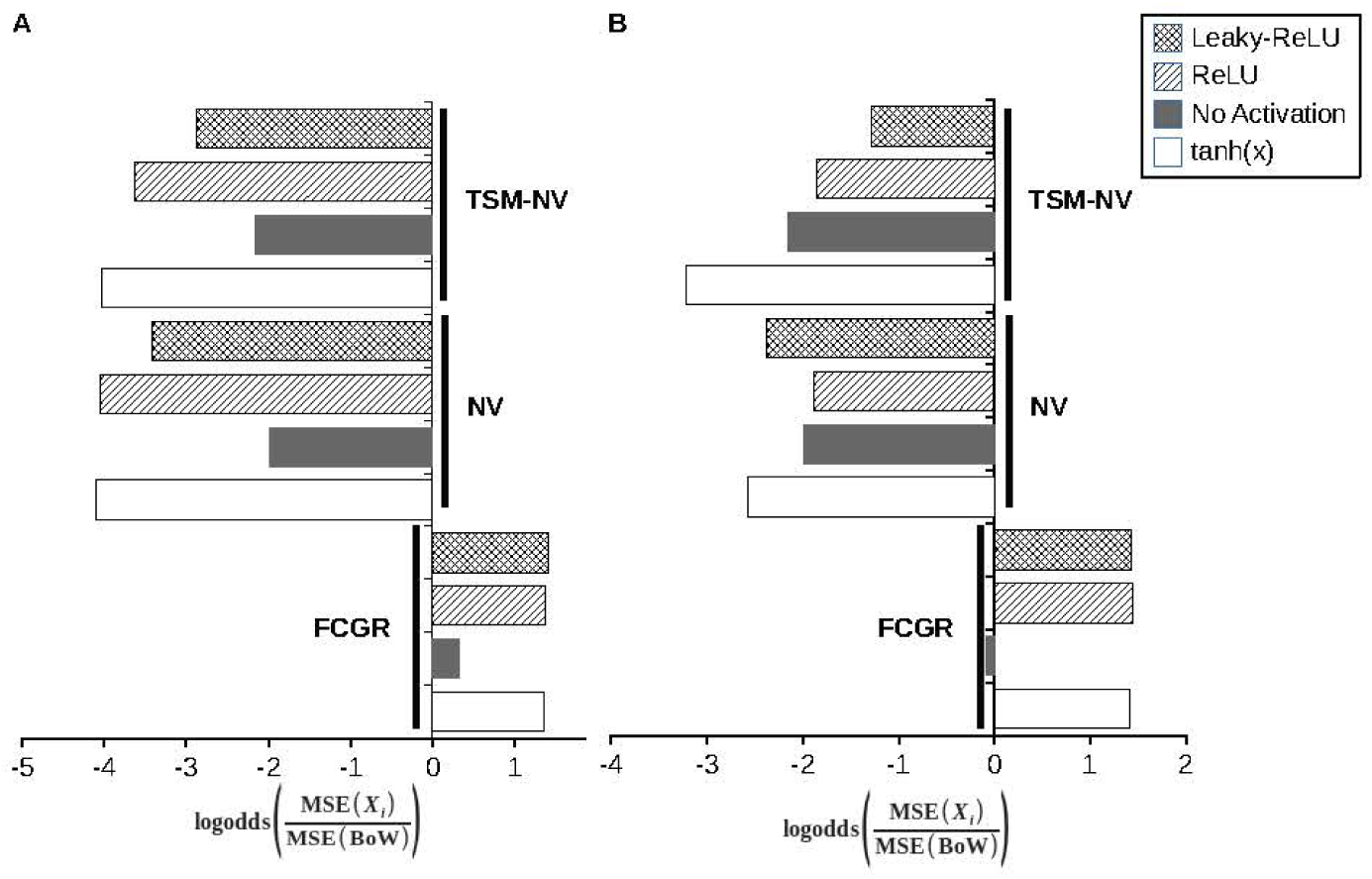
Predictive performance of MLP models trained with input data pre-processed with the evaluated methods. The horizontal bars represent the log odds for the quotient of the loss values observed in models trained with input data pre-processed with the FCGR, NV, and TSM-NV methods and the loss values for the models trained with input data pre-processed with the BoW method. In (**A**), we showed the results for models without dropout layers, and in (**B**), we show the results for models with dropout layers. The colors and hashing of the bars represent the activation functions that are described in the legend.

### Predicting percentage of identity of pairwise sequence alignments

We wonder if we could expect similar accuracies with models trained for predicting the identity percentage of a pairwise sequence alignment for two given sequences using the encoding methods proposed in this work. To address this question, we built eight MLP models (models’ ID: 009-016) with an input layer configured to accept two concatenated input vectors. We implemented this layer because we encoded the two nucleotide sequences (the aligned sequences) using the BoW, FCGR, NV and

TSM-NV methods, and they were passed to the neural network as one input layer. We also built models without dropout layers (009-012) and with dropout layers (013-016).

To evaluate the predictive performance, we employed the same methodology described above, used for testing MLP models for predicting the percentage of guanine and cytosine, but in this case, we tested the models (009-016) for predicting the percentage of identity of pairwise sequence alignments. To achieve this, we used the testing datasets to challenge these models, and we measured the log odds values as we previously described above.

In this analysis, the models weren’t as accurate as in the case of the models trained for predicting %GC. However, the log odds values obtained when the encoding methods NV and TSM-NV were used were lower than log odds values observed when the FCGR method was employed. We also observed that activation functions for this task are mandatory to make better predictions only for the NV and TSM-NV methods. Those results suggest that, despite a modest performance, training MLP neural networks with the NV and TSM-NV vectors as input data for the prediction of identity percentage given a pairwise sequence alignment, produces more accurate models compared to the models trained with input data pre-processed by the BoW and FCGR methods (**Figure 2**).

**Figure 2.**
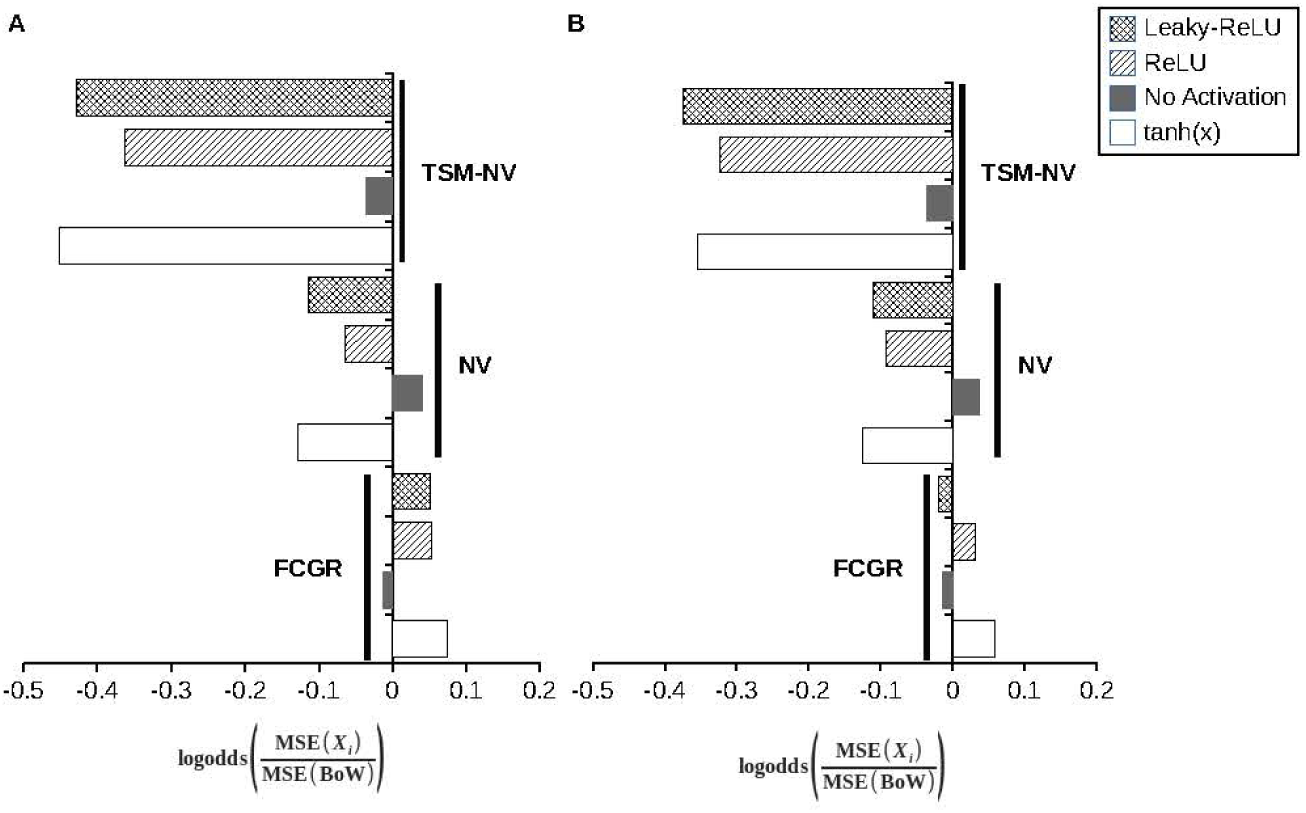
Predictive performance of the MLP models for predicting the percentage of sequence identity. The horizontal bars represent the log odds for the quotient of the loss values of models trained with input data encoded using the FCGR, NV, or TSM-NV method and the loss values for the models trained with input data encoded using the BoW method (reference). In (**A**), we showed the results for models without dropout layers, and in (**B**), we show the results for models with dropout layers. The colors and hashing of the bars represent the activation functions, which are described in the legend.

In (**Figure 3**) we show the scatter plots of the predicted and real percentage of guanine and cytosine (GC) and identity percentage of pairwise sequence alignment inferred with the best model. It is clearly observed that the prediction of guanine and cytosine percentage and the percentage of identity was more accurate in MLP models trained with the data encoded by the NV or TSM-NV methods than the MLP models trained with data encoded by the BoW and FCGR methods.

**Figure 3.**
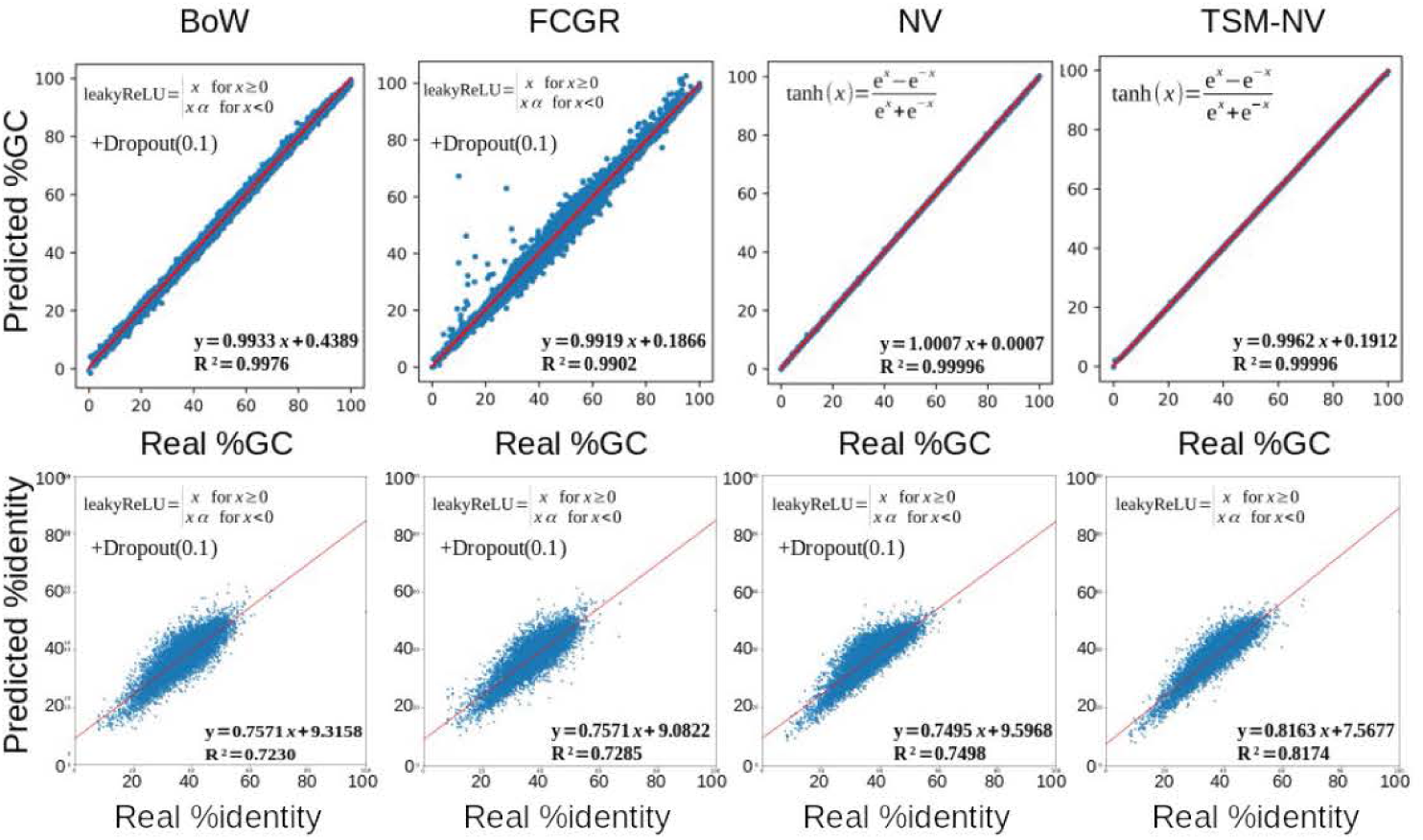
Best MLP models trained for predicting the percentage of GC and pairwise sequence identity. In this figure, we plotted the relationship between the predicted %GC and % of identity between two aligned sequences, and the real values. The determinant coefficient (R ²) and the slope of the linear model were calculated as a measure of accuracy.

### Accurate neural network classifiers were trained with data encoded by NV and TSM-NV methods

In light of these results, we asked if TSM-NV and NV encoding methods could also be suitable for training MLP models for the categorization tasks. In this case, we trained MLP models (ID 017 - 024) to predict sequence categories based on the percentage of guanine and cytosine (%GC). One group, called LOW, contained sequences with a %GC in a range of (0%-33%], another group, called MEDIUM, contained sequences with a %GC in a range of (33%-66%], and the third group, called HIGH, contained sequences with a %GC in a range of (66%-100%). As a measure of accuracy, we calculated the log odds ratio of the F1 Score as a measure of balanced accuracy between sensitivity and positive predictive value.

In (**Figure 4**), the log odds ratio of the F1Score are shown. We observed that the log odds ratio of MLP models trained with data encoded by the TSM-NV or NV methods was higher than the log odds ratio of the MLP models trained with input data pre-processed by BoW or FCGR methods, being the TSM-NV encoding method more suitable for the training models, only when leaky-ReLU and ReLU activation functions were used and no dropout layers were set. FCGR was the worst pre-processing method evaluated in this analysis because it showed the lowest log odds ratio values.

**Figure 4.**
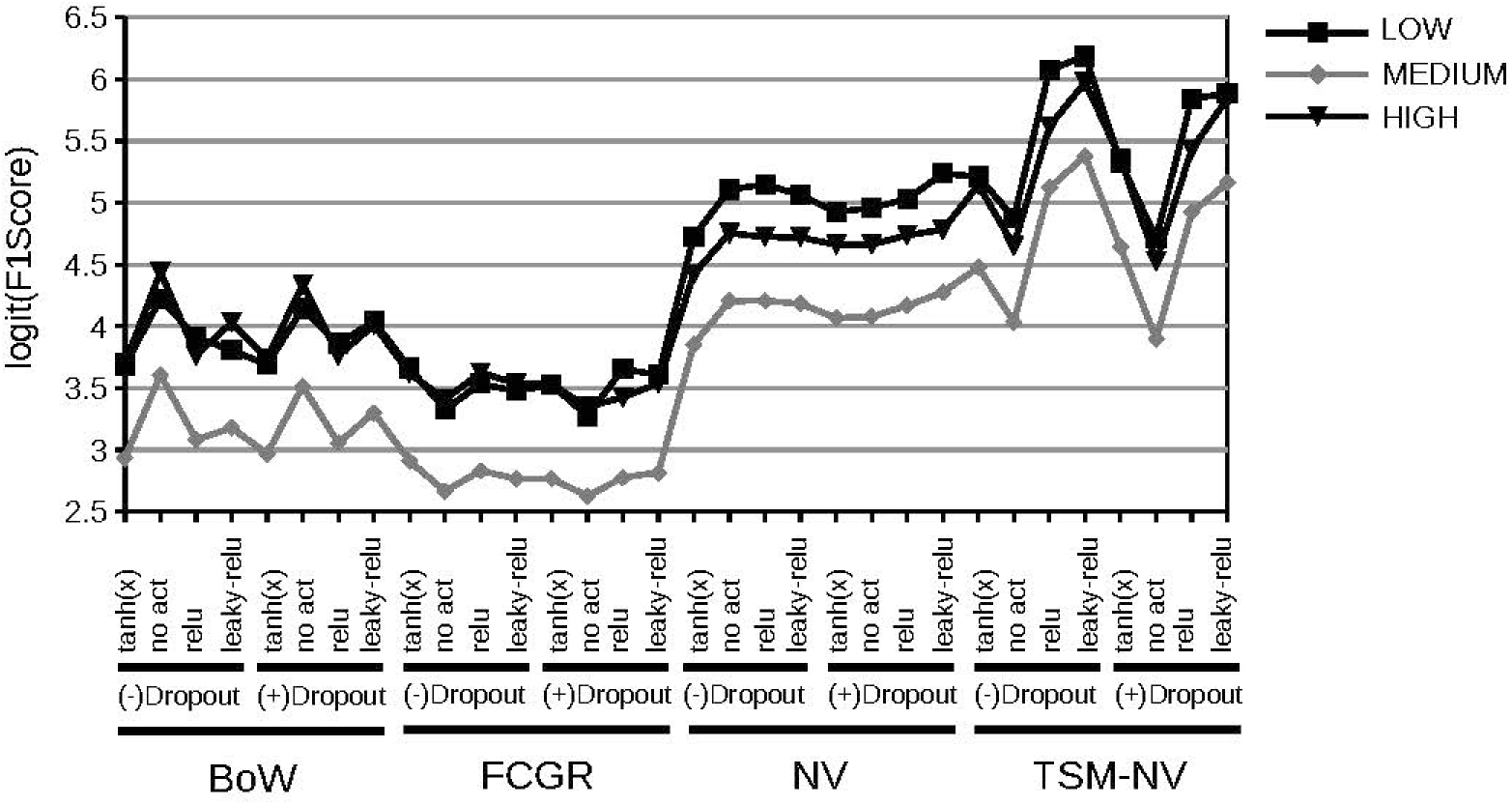
MLP models’ predictive performance in categorizing sequences based on the percentage of guanine and cytosine. MLP models were trained with input data pre-processed with the methods evaluated in this work (BoW, FCGR, NV and TSM-NV) with different activation functions (tanh, relu, leaky-relu and no activation, a.k.a. linear activation) and the use or not of dropout layers. In the abscises, we plotted the log odds ratio of F1Score/(1-F1Score).

**Figure 5.**
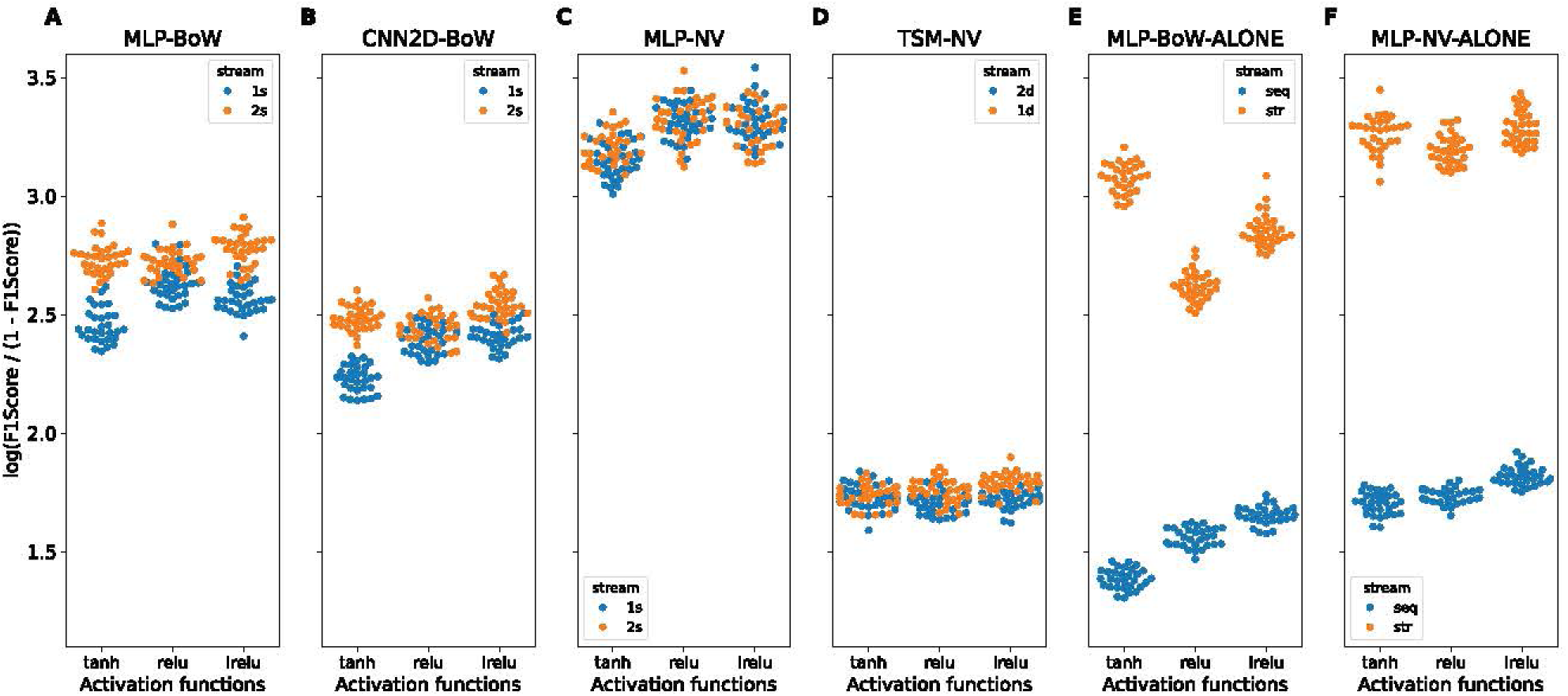
Evaluation of the artificial neural networks trained for predicting pre-miRNA sequences. In this figure we plotted the natural logarithm of the odds ratio between F1Score and (1-F1Score) observed in the evaluation of (**A**) MLP models trained with BoW vectors, (**B**) CNN2D models trained with BoW matrices, (**C**) MLP models trained with NV with covariance component, (**D**) MLP models (1d) and CNN2d (2d) models trained with flat TSM-NV vectors or TSM-NV matrices respectively, (**E**) MLP models trained with BoW vectors calculated with nucleotide sequences (seq) or sequences of the secondary structure representation (str), (**F**) MLP models trained with NV plus covariance component calculated from nucleotide sequences (seq) or secondary structure representation sequence (str). In (**A, B** and **C**), the input stream method was also tested (1s or 2s).

### Efficient training of pre-miRNA sequence classifiers using the NV method

In the previous *in silico* analysis, where computer-generated nucleotide sequence data were used during the models’ training and testing, we observed accurate performance of models trained with natural vectors with covariates and natural vectors nested within the three sequence method, as input data; these preliminary results suggest that TSM-NV and NV encoding methods efficiently represent nucleotide sequences, so we were interested if these pre-processing methods can be also suitable for the encoding of nucleotide sequences and the secondary structure representation of miRNA precursor sequences. Accordingly, we decided to carry out the training and testing of models for predicting pre-miRNA sequences of the Fabaceae plants, a challenging task that has been addressed with deep learning techniques in the past decade, but little is known about the using NV and TSMN-NV methods to encode nucleotide sequence information and it’s secondary structure for the nural network training for this purpose (Bohnsack et al., 2022). In the following analyses, we evaluated the effect of the BoW, NV, TSM-NV input data encoding methods, two neural network architectures (MLP and CNN2D), two different ways of input streaming, and three activation functions (tanh, relu, leaky-relu), on the models’ predictive performance for the classification of Fabaceae plants pre-miRNA sequences.

For these experiments, we used the nucleotide sequence (seq) and it’s secondary structure representation (str) built with the ViennaRNA package (Lorenz et al., 2011) as input data encoded by the methods we proposed in this work. We tested different ways of stream input, these were one stream (1s), where seq and str information was concatenated first and then passed to the model, and two streams (2s), where each pre-processed input (seq or str) was passed to the model by independent streams composed with 3 hidden layer of MLP or CNN2D architecture, then the output layers from these streams were concatenated and canalized to an MLP architecture with 4 hidden layers.

Consequently, we built 6 MLP models (ID 025 - 030), three of them were constructed with a one input stream (1s), and the other three with a two input stream (2s), and the activation functions hyperbolic tangent, ReLU, Leaky-ReLU (tanh, relu, leaky-relu) set between layers were varied. Then we evaluated the performance on the pre-miRNA sequence classification with the datasets described in materials and methods by sampling with replacement the testing dataset 30 times with a sample size of 5000. In (**Figure 6**), we show the natural logarithm of the <u>^f1score^</u> quotient as a measure of the classification performance for every model tested, and in (**Figure 6A**), we show the performance of the models that were trained with input data encoded with the BoW method. When the one-channel input stream (1s) was employed, we observed a little diminishing of performance only when tanh and Leaky-ReLU activation functions were set between layers, compared with the results obtained when the 2s input stream was used. We also observed better performance when ReLU and Leaky-ReLU activation functions were set between layers of the models configured with a 1s input stream. Predictive performance of MLP models configured with a 2s input stream showed no significant differences along the activation functions. Next, we speculated that reshaping the BoW vectors from one to two dimensions would form patterns that could be efficiently extracted with CNN2D models.

In (**Figure 6B**), we show the performance results for 6 CNN2D (CNN2D-BoW) models (ID 031 - 036) similarly built to the MLP models depicted above. But for this case we had to reshape the BoW vector dimension. BoW vectors for nucleotide sequences with a kmer size of 5 yielded a vector with 1024 elements, and the BoW vectors for the secondary structure representation with the same kmer size yielded a vector of 243 elements. Thus, the nucleotide sequence matrix dimension was configured to 32×32, and the secondary structure representation matrix dimension was configured to 9×27. For these models, the first 3 hidden layers were convolutional, the output was canalized to an MLP architecture with 4 hidden layers, and there were models configured with 1s and 2s input streams. We observed a very similar pattern to what we observed in the MLP-BoW experiment (**Figure 6A**), but in this case, the performance of the models was, in general, lower than the MLP-BoW models. In addition, the CNN2D-BoW models configured with 1s input stream, and with ReLU and Leaky-ReLU activation functions showed better predictive performance than the CNN2D model with tanh activation function. Models configured with 2s input stream showed no significant difference in their predictive performance (**Figure 6B**). We think the coarse way of BoW-vectors dimension reshaping produced meaningless features or deformed important one-dimensional patterns that could affect the information extraction process performed by the convolutional layers of the CNN2D models.

As we observed good performance on predicting percentage of guanine and cytosine, predicting the percentage of identity of two aligned sequences, and categorizing sequences based on guanine and citosine percentage, in early experiments with MLP models trained with natural vectors with covariates, we hypothesized that training MLP models configured with 1s input stream and 2s input stream and trained with these natural vectors, would improve the performance for the prediction of pre-miRNA sequences compared with MLP-BoW models. With this in mind, we constructed 6 MLP models (MLP-NV, ID 037 - 042) similar to the MLP-BoW experiment, but in this case we configured the input layer to receive natural vectors with covariates calculated from nucleotide sequences, and it’s secondary structure representation from training datasets. We also assessed the activation functions (tanh, relu, leaky-relu) and the input streams (1s, 2s). In (**Figure 6C**), we show the results of the predictive performance of the MLP-NV models, and we observed no significant difference in the models configured with 1s and 2s input streams but, as we expected, we observed a measurable increase in predictive performance of these models compared with the MLP-BoW models (**Figure 6C and 6A**).

Three sequences method is another technique used to represent nucleotide sequence data, it produces a matrix with a dimension of 3x*L*, where *L* stands for the nucleotide sequence length (see materials and methods section). If every vector of this matrix is transformed to natural vectors with covariates, it is possible to numerically represent the nucleotide sequence as a matrix with dimensions 3x *M* (TSM-NV matrix) where *M* represents the length of every natural vector, and we speculated that CNN2D neural networks could extract useful information to accurately predict pre-miRNA sequences. Accordingly, we used the natural vector with covariates nested within three sequence encoding method to represent the nucleotide sequences for the training of CNN2D (2d) and MLP (1d) models (ID 043 - 048). In (**Figure 6D**), we observed that the performance in predicting pre-miRNA sequences of the CNN2D and MLP models trained with the representation methods described above showed lower predictive power compared with the models of the previous MLP-BoW, CNN2D-BoW and MLP-NV experiments. The results we obtained suggest that the CNN2D and MLP models designed in this work didn’t extract useful information from the nucleotide sequences represented with the TSM-NV matrices and the flattened TSM-NV vectors.

Nevertheless, it is noteworthy that models trained with TSM-NV matrices or flattened vectors were only trained with information that came from nucleotide sequences, lacking the information source of the secondary-structure representation sequence. It is well documented the lacking of nucleotide sequence patterns between pre-miRNA and no pre-miRNA sequences and the difficult to accurately identify these precursor sequences (Meyers et al., 2008), and the observation of the well predictive performance of models trained with information that came from both nucleotide sequences and its secondary structure representation (**Figure 6A, 6B and 6C**) compared with the models trained only with nucleotide sequence information (**Figure 6D**), suggests that there is little useful information that come form nucleotide sequence, and that the well predictive power of models observed in the MLP-BoW, CNN2D-BoW and MLP-NV analyses compared with the models of TSM-NV analysis are due to the information obtained from the secondary-structure representation sequence. Therefore, we hypothesized that models trained only with information obtained from secondary-structure representation sequences will perform better than those models trained only with information that came from nucleotide sequences.

To test this hypothesis, we constructed 6 MLP models (ID 049 - 054) configured to receive BoW vectors (MLP-BoW-ALONE), three of them only received the BoW vectors calculated from nucleotide sequences (seq information), and the other three models only received the information from secondary-structure representation of the sequences (str information) (**Figure 6E**). We also built six other MLP models (ID 055 - 060) similar to the models described above, but in this case, we configured them to receive natural vectors with covariates (MLP-NV-ALONE) (**Figure 6F**), and in both analysis, the activation functions were varied. The results of these analyses agree with the predictions of the hypothesis; MLP models that were trained only with the nucleotide sequence information showed more inferior predictive performance than models trained only with the information obtained only from the secondary-structure representation sequence.

MLP-BoW models configured with tanh and Leaky-ReLU activation functions between hidden layers and trained only with the secondary-structure representation information showed a slightly significant better performance than the MLP-BoW models trained with both information sources (seq and str) and with both input stream configurations (1s and 2s). This observation raises the idea that the BoW vectors obtained from nucleotide sequence information compromised the training of MLP models that used the hyperbolic tangent (tanh) and Leaky-ReLU activation functions (**Figure 6E and 6A**). The results from these experiments, where MLP models were trained with natural vectors with covariates calculated only from the secondary-structure information, showed no significant difference compared to MLP models trained with natural vectors calculated from both information sources, nucleotide sequences and it’s secondary structure information (**Figure 6F and 6C**).

## 4. DISCUSSION

Early artificial intelligence approaches, employed in the prediction of pre-miRNA sequences or miRNA coding genes, were mainly based on Support Vector Machines (SVM), Kernel Density Estimators (KDE) and Random Forest (RF) algorithms, and several interesting methods to address the numeric representation of nucleotide sequences, secondary structure and thermodynamic properties of miRNA precursors were proposed. Sewer et al (2005) built one of the earlier SVM classifiers trained for the miRNA precursors identification that preprocesses the input data as a vector composed of 16 features extracted from the nucleotide sequence, secondary structure of the hairpin pre-miRNA. Other SMV classifiers were implemented with new strategies for feature selection, and the number of features extracted from the primary sequences, secondary structure of the hairpin and thermodynamic properties was increased. (Batuwita and Palade, 2009; Hertel and Stadler, 2006; Huang et al., 2007; Ng and Mishra, 2007; Tempel et al., 2015). Random Forest and Kernel Density Estimators were also proposed in machine learning techniques that used large feature vectors extracted from nucleotide sequences, secondary structure and thermodynamic properties. These models showed reasonable success in the pre-miRNA sequence classification (Chang et al., 2008; Cui et al., 2015; Leclercq et al., 2013). Although the performance of the models proportionally increased with the increment of the number of features and their complexity, the process of feature extraction became time-consuming and domain-expertise dependent. Feature vectors composed of local-contiguous paired information or interdependent mutual information calculated from nucleotide sequences and the secondary structure representation of miRNA precursors were implemented in SVM and RF classifiers, showing that mathematical methods applied for the numerical encoding of pre-miRNA sequences and its secondary structure can be efficiently used in machine learning (Fu et al., 2019; Jiang et al., 2007; Xue et al., 2005).

One of the earlier multi-layer perceptron models, which was trained for identifying miRNA precursor sequences, was proposed by (Gupta et al., 2012). This MLP model uses an input vector composed of 7 features, mainly extracted from the secondary structure of the hairpin, and the nucleotide sequence. This work showed that neural networks can be applied to the pre-miRNA identification. Several research works that implemented MLP models for the same purpose adopted the feature vectors previously implemented in SVM and RF classifiers as input data, adopting at the same time, the same problem of the feature extraction process (Lockuge et al., 2022; Rahman et al., 2012). Feature vectors composed of n-grams (a.k.a vectors of k-mer frequencies) were also proposed for the numerical representation of miRNA precursor nucleotide sequence and its secondary structure in MLP classifiers for the pre-miRNA prediction (Jiang et al., 2016; Montalvo-Arredondo and Juárez-Verdayes, 2025), in addition to the triplet structure sequence information (a.k.a local-contiguous paired information (Xue et al., 2005)). These encoding methods showed measurable success in the classification task (Jiang et al., 2016), suggesting that mathematical methods for the encoding of nucleotide sequence, such as n-grams and local-contiguous paired information, efficiently represent miRNA precursors.

Other neural network architectures were proposed for the pre-miRNA identification, such as Recurrent Neural Network based on the Long Short Term Memory (LSTM), and Convolutional Neural Networks (CNN). The main method used for encoding information from miRNA precursors in these neural networks was based on the one-hot encoding (Cha et al., 2021; Do et al., 2018; Georgakilas et al., 2020; Tasdelen and Sen, 2021; Zheng et al., 2019; Zheng et al., 2020). Other interesting input pre-processing methods were also proposed for these CNN and RNN models, showing descent performance, namely, word embedding (Park et al., 2016), nucleotide pairing matrices for the hairpin secondary structure representation (Do et al., 2018), pixel maps to visually represent the secondary structure and nucleotide sequence in one image (Cordero et al., 2020) and the pre-processing of input data independently by different branches or streaming channels that converge in an MLP architecture (Cha et al., 2021; Georgakilas et al., 2020).

The input data pre-processing method is one of the first steps in the training of machine learning models, and it has to extract robust and confident information from the miRNA precursor sequences and the secondary structure of the hairpin. Despite the large diversity of pre-processing methods that have been evaluated, there are other published methods for the encoding of nucleotide sequences that have been mainly used in alignment-free sequence comparison methods that have not been tested in machine learning approaches for the miRNA precursors identification (see review Bohnsack et al., 2022). Natural vector with covariance component is a nucleotide sequence encoding method that has caught our attention because it successfully represents genome sequences, and it has been used in the construction of the geometric space of the viral genomes for the correct classification of these organisms, and to understand their phylogenetic relationship. This encoding method was also used to numerically represent genomes, not only for viruses, but for bacteria, fungi and archaea species. Recently, this method was tested in neural network trained for the classification of SARS-CoV-2 and poliovirus genomes with notably success (Sun et al., 2021; Sun et al., 2022; Shi et al., 2025).

We hypothesize that natural vectors with covariance component will efficiently represent the nucleotide sequence and the hairpin secondary structure of the miRNA precursors, extracting useful information for training machine learning models applied to genomics and transcriptomics in plants or other living being species. In this work, we tested this sequence encoding method for the pre-processing of nucleotide sequences or the secondary structure representation for the training of MLP and CNN models. First, to evaluate if this method (NV) alone or nested within the Three Sequence Method (TSM-NV), could represent intrinsic properties of the nucleotide sequence, such as percentage of guanine and cytosine, and percentage of identity given an alignment of two sequences. Second, to evaluate if this encoding method (NV) alone or nested within the TSM efficiently represent the nucleotide sequence and the secondary structure of the hairpin for the classification of miRNA precursors. In this work, we observed that the NV method, alone or in combination with the TSM, successfully extracts robust and confident information from the intrinsic properties of the randomly-generated nucleotide sequences and from the miRNA precursor sequences and their hairpin secondary structure. The MLP models trained with NV showed, on average, an F1-score of 97%, 95% of sensitivity and 98% of specificity. We think these values represent a competitive performance compared with the results published in the previous works.

We also observed that in some scenarios (**Figure 6A and 6B**), the independent handling of the two input sources, namely, nucleotide sequence and hairpin secondary structure, improved the performance of the models trained with two-stream input data (2s). This trained method was also implemented by (Cha et al., 2021; Georgakilas et al., 2020) in the CNN models trained for the pre-miRNA classification and for the transcription start prediction of miRNA genes with highly success. Additionally, we observed a slight tendency to gain better performance when using the ReLU and Leaky-ReLU activation functions in classification tasks. Finally, it is noteworthy that the negative effect of training the MLP models with BoW vectors (**Figure 6E**) and the little positive effect of training MLP models with NV (**Figure 6F**) extracted only from nucleotide sequences, these results suggest that there are no real patterns that permit discerning between pre-miRNA and no pre-miRNA sequences.

In summary, the results of this work suggest that natural vectors with covariates could be a streamlined option for the pre-processing of the input data for the neural network training. Although natural vectors with covariance component were successfully tested in MLP and in some CNN architectures, the current approach of this work is straightforward, so we think it might benefit from further exploration of other neural network architectures, advanced mathematical analysis and different training datasets to exploit the potential of the natural vectors.

## REFERENCES

Ayubov, M. S., Mirzakhmedov, M. H., Sripathi, V. R., Buriev, Z. T., Ubaydullaeva, K. A., Usmonov, D. E., … Abdurakhmonov, I. Y. (2019). Role of MicroRNAs and small RNAs in regulation of developmental processes and agronomic traits in Gossypium species. Genomics, 111(5), 1018–1025. doi: 10.1016/j.ygeno.2018.07.012

Baez-Mora, G., Rueda-Torres, L., Gonzalez-Morgado, A., Valle-Harris, C., & Díaz-Nava, C. (2023). Socioeconomic characteristics of the broad bean (Vicia faba L.)(Fabaceae) production in the northeastern region of the State of Puebla, Mexico. Agro Productividad. doi: 10.32854/agrop.v16i7.2493

Batuwita, R., & Palade, V. (2009). microPred: effective classification of pre-miRNAs for human miRNA gene prediction. Bioinformatics, 25(8), 989–995. doi: 10.1093/bioinformatics/btp107

Bohnsack, K. S., Kaden, M., Abel, J., & Villmann, T. (2022). Alignment-free sequence comparison: A systematic survey from a machine learning perspective. IEEE/ACM Transactions on Computational Biology and Bioinformatics, 20(1), 119–135. doi: 10.1109/TCBB.2022.3140873

Brant, E. J., & Budak, H. (2018). Plant small non-coding RNAs and their roles in biotic stresses. Frontiers in Plant Science, 9, 1038. doi: 10.3389/fpls.2018.01038

Budak, H., Kantar, M., Bulut, R., & Akpinar, B. A. (2015). Stress responsive miRNAs and isomiRs in cereals. Plant Science, 235, 1–13. doi: 10.1016/j.plantsci.2015.02.008

Centeno-González, N. K., Martínez-Cabrera, H. I., Porras-Múzquiz, H., & Estrada-Ruiz, E. (2021). Late Campanian fossil of a legume fruit supports Mexico as a center of Fabaceae radiation. Communications Biology, 4(1), 41. doi: 10.1038/s42003-020-01533-9

Cha, M., Zheng, H., Talukder, A., Barham, C., Li, X., & Hu, H. (2021). A two-stream convolutional neural network for microRNA transcription start site feature integration and identification. Scientific Reports, 11(1), 5625. doi: 10.1038/s41598-021-85173-x

Chang, D. T. H., Wang, C. C., & Chen, J. W. (2008). Using a kernel density estimation based classifier to predict species-specific microRNA precursors. BMC Bioinformatics, 9(Suppl 12), S2. doi: 10.1186/1471-2105-9-S12-S2

Cordero, J., Menkovski, V., & Allmer, J. (2020). Detection of pre-microRNA with convolutional neural networks. BioRxiv, 840579. doi: 10.1101/840579

Cui, H., Zhai, J., & Ma, C. (2015). miRLocator: machine learning-based prediction of mature microRNAs within plant pre-miRNA sequences. PLoS One, 10(11), e0142753. doi: 10.1371/journal.pone.0142753

Delgado-Salinas, A., Torres-Colín, L., Luna-Cavazos, M., & Bye, R. (2021). Diversity of useful Mexican legumes: analyses of herbarium specimen records. Diversity, 13(6), 267. doi: 10.3390/d13060267

Do, B. T., Golkov, V., Gürél, G. E., & Cremers, D. (2018). Precursor microRNA identification using deep convolutional neural networks. BioRxiv, 414656.doi: 10.1101/414656

Dong, Q., Hu, B., & Zhang, C. (2022). microRNAs and their roles in plant development. Frontiers in plant science, 13, 824240. doi: 10.3389/fpls.2022.824240

Estrada-Castillón, E., Villarreal-Quintanilla, J. Á., Cuéllar-Rodríguez, G., Encina-Domínguez, J. A., Martínez-Ávalos, J. G., Mora-Olivo, A., & Sánchez-Salas, J. (2024). The Fabaceae in Northeastern Mexico (Subfamily Caesalpinioideae, Mimosoideae Clade, Tribes Mimoseae, Acacieae, and Ingeae). Plants, 13(3), 403. doi: 10.3390/plants13030403

Feng, S., Xu, Y., Guo, C., Zheng, J., Zhou, B., Zhang, Y., … Wang, H. (2016). Modulation of miR156 to identify traits associated with vegetative phase change in tobacco (Nicotiana tabacum). Journal of Experimental Botany, 67(5), 1493–1504. doi: 10.1093/jxb/erv551

Fu, X., Zhu, W., Cai, L., Liao, B., Peng, L., Chen, Y., & Yang, J. (2019). Improved pre-miRNAs identification through mutual information of pre-miRNA sequences and structures. Frontiers in Genetics, 10, 119. doi: 10.3389/fgene.2019.00119

Gangadhar, B. H., Venkidasamy, B., Samynathan, R., Saranya, B., Chung, I. M., & Thiruvengadam, M. (2021). Overview of miRNA biogenesis and applications in plants. Biologia, 76(8), 2309–2327. doi: 10.1007/s11756-021-00763-4

Georgakilas, G. K., Grioni, A., Liakos, K. G., Chalupova, E., Plessas, F. C., & Alexiou, P. (2020). Multi-branch convolutional neural network for identification of small non-coding RNA genomic loci. Scientific Reports, 10(1), 9486. doi: 10.1038/s41598-020-66454-3

Guo, Z., Kuang, Z., Zhao, Y., Deng, Y., He, H., Wan, M., … & Li, L. (2022). PmiREN2. 0: from data annotation to functional exploration of plant microRNAs. Nucleic Acids Research, 50(D1), D1475–D1482. doi: 10.1093/nar/gkab811

Gupta, M. K., Agarwal, K., Prakash, N., & Singh, D. B. (2012). Prediction of miRNA in HIV-1 genome and its targets through artificial neural network: a bioinformatics approach. Network Modeling Analysis in Health Informatics and Bioinformatics, 1(4), 141–151. doi: 10.1007/s13721-012-0017-3

Hertel, J., & Stadler, P. F. (2006). Hairpins in a Haystack: recognizing microRNA precursors in comparative genomics data. Bioinformatics, 22(14), e197–e202. doi: 10.1093/bioinformatics/btl257

Huang, T. H., Fan, B., Rothschild, M. F., Hu, Z. L., Li, K., & Zhao, S. H. (2007). MiRFinder: an improved approach and software implementation for genome-wide fast microRNA precursor scans. BMC Bioinformatics, 8, 341. doi: 10.1186/1471-2105-8-341

Jiang, L., Zhang, J., Xuan, P., & Zou, Q. (2016). BP Neural Network Could Help Improve Pre-miRNA Identification in Various Species. BioMed Research International, 2016, 9565689. doi: 10.1155/2016/9565689

Jiang, P., Wu, H., Wang, W., Ma, W., Sun, X., & Lu, Z. (2007). MiPred: classification of real and pseudo microRNA precursors using random forest prediction model with combined features. Nucleic Acids Research, 35(suppl_2), W339-W344. doi: 10.1093/nar/gkm368

Kozomara, A., & Griffiths-Jones, S. (2011). miRBase: integrating microRNA annotation and deep-sequencing data. Nucleic acids research, 39(suppl_1), D152-D157. doi: 10.1093/nar/gkq1027

Leclercq, M., Diallo, A. B., & Blanchette, M. (2013). Computational prediction of the localization of microRNAs within their pre-miRNA. Nucleic Acids Research, 41(15), 7200–7211. doi: 10.1093/nar/gkt466

Li, S., Castillo-González, C., Yu, B., & Zhang, X. (2017). The functions of plant small RNAs in development and in stress responses. The Plant Journal, 90(4), 654–670. doi: 10.1111/tpj.13444

Löchel, H. F., & Heider, D. (2021). Chaos game representation and its applications in bioinformatics. Computational and Structural Biotechnology Journal, 19, 6263–6271. doi: 10.1016/j.csbj.2021.11.008

Lokuge, S., Jayasundara, S., Ihalagedara, P., Kahanda, I., & Herath, D. (2022). miRNAFinder: A comprehensive web resource for plant Pre-microRNA classification. Biosystems, 215, 104662. doi: 10.1016/j.biosystems.2022.104662

Lorenz, R., Bernhart, S. H., Höner zu Siederdissen, C., Tafer, H., Flamm, C., Stadler, P. F., & Hofacker, I. L. (2011). ViennaRNA Package 2.0. Algorithms for Molecular Biology, 6(1), 26. doi: 10.1186/1748-7188-6-26

Martinc, M., & Pollak, S. (2019). Combining n-grams and deep convolutional features for language variety classification. Natural Language Engineering, 25(5), 607–632. doi: 10.1017/S1351324919000299

Meyers, B. C., Axtell, M. J., Bartel, B., Bartel, D. P., Baulcombe, D., Bowman, J. L., … & others. (2008). Criteria for annotation of plant MicroRNAs. The Plant Cell, 20(12), 3186–3190. doi: 10.1105/tpc.108.064311

Ng, K. L. S., & Mishra, S. K. (2007). De novo SVM classification of precursor microRNAs from genomic pseudo hairpins using global and intrinsic folding measures. Bioinformatics, 23(11), 1321–1330. doi: 10.1093/bioinformatics/btm026

Ouyang, S., Park, G., Atamian, H. S., Han, C. S., Stajich, J. E., Kaloshian, I., & Borkovich, K. A. (2014). MicroRNAs suppress NB domain genes in tomato that confer resistance to Fusarium oxysporum. PLoS pathogens, 10(10), e1004464. doi: 10.1371/journal.ppat.1004464

Park, S., Min, S., Choi, H. S., & Yoon, S. (2016). deepMiRGene: Deep neural network based precursor microRNA prediction. arXiv preprint arXiv:1605.00017.doi: 10.48550/arXiv.1605.00017

Rahman, M. E., Islam, R., Islam, S., Mondal, S. I., & Amin, M. R. (2012). MiRANN: A reliable approach for improved classification of precursor microRNA using Artificial Neural Network model. Genomics, 99(4), 189–194. doi: 10.1016/j.ygeno.2012.02.001

Sewer, A., Paul, N., Landgraf, P., Aravin, A., Pfeffer, S., Brownstein, M. J., … & Zavolan, M. (2005). Identification of clustered microRNAs using an ab initio prediction method. BMC Bioinformatics, 6(1), 267. doi: 10.1186/1471-2105-6-267

Shavanov, M. V. (2021). The role of food crops within the Poaceae and Fabaceae families as nutritional plants. IOP Conference Series: Earth and Environmental Science, 624(1), 012111. doi: 10.1088/1755-1315/624/1/012111

Shi, X., Kang, J., Sun, N., Zhao, X., & Yau, S. S. T. (2025). Multi-Perspective Natural Vector: A Novel Method for Viral Sequence Feature Extraction. Journal of Computational Biology. doi: 10.1177/15578666251391211

Shriram, V., Kumar, V., Devarumath, R. M., Khare, T. S., & Wani, S. H. (2016). MicroRNAs as potential targets for abiotic stress tolerance in plants. Frontiers in Plant Science, 7, 817. doi: 10.3389/fpls.2016.00817

Smith, T. F., & Waterman, M. S. (1981). Identification of common molecular subsequences. Journal of Molecular Biology, 147(1), 195–197.

Song, X., Li, Y., Cao, X., & Qi, Y. (2019). MicroRNAs and their regulatory roles in plant--environment interactions. Annual review of plant biology, 70(1), 489–525. doi: 10.1146/annurev-arplant-050718-100334

Su, Y., Li, H. G., Wang, Y., Li, S., Wang, H. L., Yu, L., … & others. (2018). Poplar miR472a targeting NBS-LRRs is involved in effective defence against the necrotrophic fungus Cytospora chrysosperma. Journal of Experimental Botany, 69(22), 5519–5530. doi: 10.1093/jxb/ery304

Sun, N., Pei, S., He, L., Yin, C., He, R. L., & Yau, S. S. T. (2021). Geometric construction of viral genome space and its applications. Computational and Structural Biotechnology Journal, 19, 4226–4234. doi: 10.1016/j.csbj.2021.07.028

Sun, N., Zhao, X., & Yau, S. S. T. (2022). An efficient numerical representation of genome sequence: natural vector with covariance component. PeerJ, 10, e13544. doi: 10.7717/peerj.13544

Tasdelen, A., & Sen, B. (2021). A hybrid CNN-LSTM model for pre-miRNA classification. Scientific Reports, 11(1), 14125. doi: 10.1038/s41598-021-93656-0

Tempel, S., Zerath, B., Zehraoui, F., Tahi, F., & others. (2015). miRBoost: boosting support vector machines for microRNA precursor classification. RNA, 21(5), 775–785. doi: 10.1261/rna.043612.113

Trevethan, R. (2017). Sensitivity, specificity, and predictive values: foundations, pliabilities, and pitfalls in research and practice. Frontiers in Public Health, 5, 307. doi: 10.3389/fpubh.2017.00307

Wu, G., Park, M. Y., Conway, S. R., Wang, J.-W., Weigel, D., & Poethig, R. S. (2009). The sequential action of miR156 and miR172 regulates developmental timing in *Arabidopsis*. Cell, 138(4), 750–759. doi: 10.1016/j.cell.2009.06.031

Xue, C., Li, F., He, T., Liu, G. P., Li, Y., & Zhang, X. (2005). Classification of real and pseudo microRNA precursors using local structure-sequence features and support vector machine. BMC Bioinformatics, 6(1), 310. doi: 10.1186/1471-2105-6-310

Yang, L., Mu, X., Liu, C., Cai, J., Shi, K., Zhu, W., & Yang, Q. (2015). Overexpression of potato miR482e enhanced plant sensitivity to *Verticillium dahliae* infection. Journal of Integrative Plant Biology, 57(12), 1078–1088. doi: 10.1111/jipb.12348

Yang, X., Zhang, L., Yang, Y., Schmid, M., & Wang, Y. (2021). miRNA mediated regulation and interaction between plants and pathogens. International Journal of Molecular Sciences, 22(6), 2913. doi: 10.3390/ijms22062913

Zhang, F., Yang, J., Zhang, N., Wu, J., & Si, H. (2022a). Roles of microRNAs in abiotic stress response and characteristics regulation of plant. Frontiers in Plant Science, 13, 919243. doi: 10.3389/fpls.2022.919243

Zhang, L., Xiang, Y., Chen, S., Shi, M., Jiang, X., He, Z., & Gao, S. (2022b). Mechanisms of microRNA biogenesis and stability control in plants. Frontiers in Plant Science, 13, 844149. doi: 10.3389/fpls.2022.844149

Zhang, Y., Huang, J., Xie, F., Huang, Q., Jiao, H., & Cheng, W. (2024). Identification of plant microRNAs using convolutional neural network. Frontiers in Plant Science, 15, 1330854. doi: 10.3389/fpls.2024.1330854

Zheng, X., Fu, X., Wang, K., & Wang, M. (2020). Deep neural networks for human microRNA precursor detection. BMC Bioinformatics, 21(1), 17. doi: 10.1186/s12859-020-3339-7

Zheng, X., Xu, S., Zhang, Y., & Huang, X. (2019). Nucleotide-level convolutional neural networks for pre-miRNA classification. Scientific Reports, 9(1), 628. doi: 10.1038/s41598-018-36946-4

Zhu, Q. H., Fan, L., Liu, Y., Xu, H., Llewellyn, D., & Wilson, I. (2013). miR482 regulation of NBS-LRR defense genes during fungal pathogen infection in cotton. PloS one, 8(12), e84390. doi: 10.1371/journal.pone.0084390

